# A computational model reveals that spatial localization of cancer stem cells increases radioresistance in tumorspheres

**DOI:** 10.64898/2026.05.08.723756

**Authors:** Jerónimo Fotinós, Carlos A. Condat, Lucas Barberis

## Abstract

Cancer stem cells (CSCs) exhibit increased resistance to radiotherapy, contributing to tumor recurrence and progression. While CSCs are known for their intrinsic resistance, the role of their spatial organization remains poorly understood. We extend a computational model of tumorsphere growth to investigate how the spatial distribution of CSCs influences radiation response. The model explicitly tracks cell lineages and spatial positions, revealing a preferential accumulation of CSCs in the spheroid interior. Because radiosensitivity increases with oxygen availability, and oxygen levels are lowest in the tumor core, this spatial organization confers a protective advantage to the CSC population. We find that this effect is negligible in small, well-oxygenated tumorspheres but becomes pronounced as growth leads to the emergence of hypoxic regions. To isolate the role of spatial structure, we compare these results with control simulations in which CSC positions are randomly reassigned. In these synthetic configurations, CSC survival under irradiation is markedly reduced, demonstrating that spatial localization is a key determinant of radioresistance. This effect persists even after the onset of central necrosis, suggesting that the “spatial niche” of CSCs is a critical target for improving therapeutic outcomes.

**Author Summary:** Cancer stem cells are known to survive radiotherapy better than other cancer cells, often leading to tumor recurrence. While this resistance is usually attributed to intrinsic biological differences between cells, in this study we show that their physical location within the tumor plays a critical and previously underestimated role. We developed a three-dimensional computer model that simulates the growth of a tumorsphere from a single cancer stem cell. Because oxygen levels influence how sensitive cells are to radiation, our model tracks the position of each cell and calculates the oxygen distribution. We found that cancer stem cells naturally accumulate in the poorly oxygenated spheroid core, where radiation is less effective. To confirm that this location directly causes their survival advantage, we performed a “digital experiment“: We artificially redistributed the same cells randomly throughout the tumorsphere before applying simulated radiation. In this random configuration, cancer stem cell survival dropped significantly. Our results show that radioresistance is not only an intrinsic cell property, but also a consequence of the spatial structure of the tumor. This finding suggests that future therapies could be improved by targeting not only the stem cells themselves, but also the protective hypoxic niches where they reside.

## 1. Introduction

The development and progression of many solid tumors are determined by the presence of cancer stem cells, cells that may self-renew and differentiate, giving rise to the differentiated cancer cells (DCCs) that make up the bulk of the tumor [1–3]. Tumor recurrence and metastasis have been attributed to the ability of CSCs to resist cytotoxic drugs and radiotherapy [2, 4–9]. The prevailing view is that surviving CSCs are critical drivers of tumor relapse and metastatic spread. Consequently, effective therapeutic strategies must specifically target CSCs or their supportive microenvironments. Achieving this goal requires identifying the spatial distribution of CSCs within tumors and understanding the mechanisms underlying their resistance to therapy. Three key features are known to contribute to the exceptional radioresistance of CSCs: low levels of reactive oxygen species (ROS), enhanced DNA damage repair capacity, and quiescence [10–12]. In this work, we identify a fourth contributing factor to CSC radioresistance: their spatial localization within the tumor. Specifically, we show that CSCs accumulate in hypoxic (poorly oxygenated) regions, which renders them less sensitive to radiotherapy. Our results demonstrate that the resistance of CSCs to radiation is strongly influenced by their preferential positioning away from the surface.

The first step is to determine the spatial distribution of CSCs prior to the therapy. We focus on tumorspheres, spheroids grown in suspension from a single cancer stem cell [13–21]. In a recent experiment [22], tumorspheres enriched in CSCs were cultivated, and the immunofluorescent detection of the stemness marker SOX2 was carried out using confocal microscopy. An image processing method was then implemented to reconstruct the number and location of the CSCs in the spheroids; this analysis revealed that, consistent with the predictions of a two-dimensional model [23], CSCs accumulate in the interior of the spheroid. However, the findings of Ref. [23] do not readily generalize to three dimensions, where the larger number of available growth paths could a priori substantially alter cell distribution patterns. A realistic description of CSC-driven tumor growth therefore requires explicitly three-dimensional modelling.

We also need to describe how the local oxygen availability conditions cell survival. It has been known for many years that hypoxic tumor cells are less sensitive to radiation than well-oxygenated ones [24]. In fact, reoxygenation of tumor cells may increase tumor control [25]. This oxygen enhancement of the radiation effects is attributed to the oxygen-induced fixation of DNA damage [26]. In embryonic stem cells hypoxia is negligible for aggregates with a radius smaller than 100 μm, but it becomes an important factor for aggregates larger than 300 μm [27]. Of course, these numbers depend on the cell line, the environment, and on the spheroid tortuosity and porosity. A mathematical model to describe the role of oxygen in avascular tumor growth was developed in Refs. [26, 28] and used by Brüningk and coworkers to investigate spheroid response to radiation therapy [29]. Here we will follow Frieboes and co-workers and assume that oxygen consumption grows linearly with the oxygen concentration, i.e., that consumption is far from saturation [30]. We will also assume that the probability of cell killing is proportional to the absorption of oxygen. This probability will be taken to be the same for CSCs and DCCs, so that any differences we find in phenotype survival can be solely attributed to the cell location at the time the radiotherapy is applied.

Detailed mathematical models of the early phases of CSC-driven tumor growth and the effects of radiation have been developed over the last 15 years [31–35]. Among their findings, is the observation that radiation enriches the CSC population, potentially leading to accelerated tumor repopulation, a phenomenon sometimes referred to as the radioresistance paradox [35]. While these models, intended for in vivo tumors, predict that CSCs should be predominantly concentrated in the tumor interior, the specific contribution of spatial organization to treatment resistance has not been fully isolated. We note that a phenomenon similar to the radioresistance paradox has also been described in the context of chemotherapy [36]. Interestingly, Lagzian et al. recently proposed an agent-based model integrated with reinforcement learning (Q-learning) to simulate radiotherapy in a two-dimensional tumor initiated from a single CSC [37]. In their model, oxygen is allowed to diffuse from microchannels perpendicular to the tissue, generating localized hypoxic zones that limit proliferation and promote treatment resistance. Although this approach enables high-resolution simulations of tumor development and radioresistance in two-dimensions, it does not isolate the effect of CSC spatial location on survival.

Since our focus is to simulate experimental radiation on tumorspheres, which lack long-term dynamics, we adopt a deliberately simplified model. This approach allows us to isolate and examine the relationship between the spatial distribution of cancer stem cells and radioresistance, without introducing additional effects that are nonessential at early times and could obscure our primary objective.

Our working hypothesis is that an important reason for the observed survival of a higher fraction of CSCs than DCCs is the concentration of CSCs in the most hypoxic region of the tumorsphere. In this paper we verify the validity of this hypothesis using a computational growth model. We begin by presenting simulation results that describe the evolution of the CSC distribution in a growing tumorsphere, and then analyze the effect of a radiotherapy session on the CSC and DCC populations. We then compare our results with those obtained when spheroids containing random distributions of both subpopulations are irradiated. Finally, the impact of the application of radiotherapy after the emergence of a central necrosis is considered in detail.

## 2. Methods

### 2.1. Main Assumptions

In a standard experimental assay, it is possible to determine the population doubling time (PDT) of the total cell population. Since this measurement cannot discriminate between the growth rates of CSCs and DCCs, we will assume the same intrinsic growth rate for both populations [19]. CSC division is modeled as a stochastic process with three possible outcomes: symmetric self-renewal, yielding two CSCs with probability *p*_*s*_, symmetric differentiation, yielding two DCCs with probability *p*_*d*_, and asymmetric division, yielding one CSC and one DCC with probability *p*_*a*_ = 1 − *p*_*s*_ − *p*_*d*_. The self-replication probability *p*_*s*_ is small in homeostasis and in most common culture media. Nevertheless, CSCs can be forced to self-renew by changing their microenvironmental conditions, e.g., by using specific growth factors that inhibit differentiation. In such conditions, *p*_*s*_ can be close to unity, leading to CSC-enriched tumorspheres [38, 39]. Thus, *p*_*s*_ is the key parameter regulating the CSC fraction in simulations, capturing the effect of growth factors in tumorsphere assays. It is important to note that increasing *p*_*d*_ leads to less favorable conditions for CSC expansion. In fact, the probability of finding CSCs at the periphery (where they are more exposed to radiation) is minimized by choosing *p*_*d*_ = 0. We will therefore neglect *p*_*d*_, as this choice represents the most conservative scenario for our hypothesis while also simplifying the analysis.

Computationally, we start from a single CSC seed and, consistent with the observations of Brú et al. [40], and of Enderling et al. [31], allow it and its progeny to either duplicate or generate DCCs provided there is sufficient space for the newly produced cells. The resulting spheroids contain a core of stem and non-stem quiescent cancer cells, unable to proliferate, and a rim of proliferative cells.

### 2.2. Overview of the Computational Model

We study a lattice-free, constrained stochastic proliferation process coupled to oxygen diffusion. We use a tailored Python 3 object-oriented code built as a package, with cells in two phenotypes: CSCs and DCCs. Cells are modeled as hard spheres of radius *r*_0_. Each cell can be either in a proliferative or a quiescent state during growth, and change to a killed state after radiotherapy.

We start the simulation with a single active CSC and allow it to divide. Later, at each time step, all proliferative cells try to divide by generating candidate positions uniformly at random on a spherical surface of radius 2*r*_0_ and attempting to place the daughter cell at one of these positions. A division attempt fails if the proposed position overlaps with a preexisting cell. After *M* = 50 failed attempts, the original cell becomes quiescent; this number corresponds to the minimum needed avoid statistically significant changes in the outcome of the realizations. If the division attempt is successful and the parent cell is a CSC, it either self-renews with probability *p*_*s*_ or generates one CSC and one DCC with probability 1 − *p*_*s*_. If the parent and daughter cells belong to different phenotypes (one CSC and one DCC), we exchange the positions of the original CSC and the new DCC with probability 1/2. For DCCs, division produces only DCC offspring, rendering such exchanges inconsequential. Cell updates are made in random order; once all proliferative cells are requested to duplicate, independently of the success of their attempts, we say that a one-day-long time step has been performed. This defines a PDT of one day, consistent with typical growth rates and facilitating interpretation. Further details on the structure of the simulation algorithm are provided in the Supporting Information.

### 2.3. 0_2_ Uptake

Since we will assume that cell sensitivity to radiation is controlled by oxygen absorption, we need to know the oxygen distribution inside the tumorsphere. For small tumorspheres, we could assume that the oxygen distribution is uniform, but for larger tumorspheres, we must compute it using the diffusion equation, obtaining a concentration that depends on the distance from the spheroid center. The *O*_2_concentration is rescaled in such a way that it takes the value of unity at the spheroid edge and may thus be compared to a probability density.

The oxygen distribution depends on the rate of oxygen uptake. Various forms have been proposed for the dependence of oxygen uptake on oxygen concentration. Some authors use zero-order kinetics (absorption rate independent of local concentration) [24, 41, 42], while Frieboes and co-workers use first-order kinetics (oxygen uptake proportional to the local concentration) [30]. Other authors use saturation-type kinetics of the Michaelis-Menten [27] or exponential [43] forms. Since we are interested in considering regions with low oxygen concentrations, for which concentration-independent uptake is not a good approximation, we choose first-order kinetics. The oxygen distribution in small tumorspheres is approximately constant, but in medium-sized or large spheroids, the concentration decreases substantially as we move away from the surface. We thus need to consider oxygen diffusion and uptake inside the spheroid. Since oxygen diffusion occurs over much shorter timescales than cell replication, the oxygen distribution *c*(*r*) can be assumed to satisfy the steady state diffusion equation at all times,

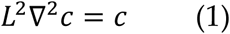

where *L* is the diffusion length,

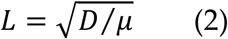

Here *D* is the oxygen diffusion coefficient and μ is the oxygen absorption rate, which is assumed to be the same for CSCs and DCCs. As mentioned before, we fix the oxygen concentration at the tumorsphere boundary to unity, i.e., *c*(*r* = *R*(*t*)) = 1, with *R*(*t*) being the instantaneous tumorsphere radius. Since the tumorsphere maintains an approximately spherical geometry throughout its evolution, operationally we define *R*(*t*)) as the average radial distance of all non-quiescent cells from the spheroid center. This definition avoids fluctuations due to isolated cells and provides a stable estimate of the effective spheroid radius. The second boundary condition is (*dc*⁄*dr*)_*r*=0_ = 0.

The value of the diffusion length may be found by considering the thickness of the viable rim. In this work, we take *L* = 100 *μm*, a representative physiological value [30]. The derived boundary value problem is solved numerically by means of a 4th-order collocation algorithm, as implemented by scipy.integrate.solve_bvp (see also [44]), and using a mesh of 50,000 nodes, which ensures high precision even for large spheroids.

### 2.4. Implementation of Therapy

We implement a phenomenological model of radiotherapy based on the Oxygen Effect, where the presence of molecular oxygen enhances the lethal effects of ionizing radiation. Cells are assumed to be killed with probability *c*(*r*), where *c*(*r*) is the concentration of *O*_2_ at a distance *r* from the spheroid center. This linear dependence is a phenomenological approximation introduced to isolate spatial effects of oxygen heterogeneity, independently of dose-response nonlinearities. We simulate the application of a single, spatially uniform radiation dose. The fraction of cells (CSCs and DCCs) killed is assumed to depend only on the local oxygen consumption rate and the applied dose, but not on the cell phenotype. Since the dose *D* is applied uniformly, we may factor out its dependence on *D* and assume that the total concentration *S*(*r*) of surviving cells is governed solely by the oxygen profile. Accordingly, the expected concentration of surviving cells satisfies,

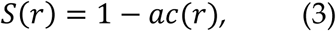

where *a* is a sensitivity coefficient. For the purpose of this study, we set *a* = 1 to represent a scenario where cells in fully oxygenated regions (*c* = 1) reach the maximum threshold of radiosensitivity, while cells in purely anoxic regions (*c* = 0) are fully protected.

In each simulation run, we select a number *ξ* at random between 0 and 1. If *ξ* > *c*(*r*), the cell survives; otherwise, it dies immediately. Finally, we group the numbers of total (*t*), surviving (*s*), and killed (*k*) cells of each phenotype (CSCs and DCCs) into spherical shells of thickness 10 *μm*, corresponding approximately to a single cell diameter. We use these data to generate histograms and measure the required fractions. Since our primary interest lies in the effect of the spatial distribution of the CSCs on radioresistance, the overall dependence on radiation dose is not central to our analysis and will not be considered further.

We emphasize that radiation is assumed to affect CSCs and DCCs in exactly the same way. To isolate the impact of the spatial distribution of CSCs, we deliberately do not attribute any intrinsic resistance to therapy to CSCs compared to DCCs. It is also convenient to describe the cell distributions inside the tumorspheres using histograms, which average the results over 32 realizations, and whose bins, of width *a* = 10 *μm*, represent the probability of finding a cell between *r* and *r* + *a*, with *r* being the distance to the spheroid center. Smooth probability density functions are then obtained through a kernel density estimation, providing continuous approximations of the spatial distributions for each cell type.

### 2.5. Random CSC Distributions

The non-uniform spatial distributions of CSCs we obtain are a direct consequence of the choices that CSCs make when they proliferate, and of space constraints. To address the protective nature of the CSC location, we compare the effect of radiotherapy on tumorspheres generated with our model with its effect on spheroids containing random distributions of CSCs. The latter were built by first taking the tumorspheres described in previous sections and removing the labels that distinguish between cell phenotypes. Then, for each of these “plain” tumorspheres, we randomly assign the CSC phenotype to the cells in such a way that they end up with the same number of CSCs as the original spheroids. Finally, we apply the therapy and collect the distributions of surviving and killed cells in each phenotype as described in previous sections. We repeat this redistribution and therapy process 50 times on each plain tumorsphere to increase the statistical confidence of this study. Comparing the outcomes of radiotherapy applied to model-generated versus random CSC distributions allows us to isolate the impact of spatial geometry on effective CSC radioresistance.

### 2.6. Tumorspheres with Necrotic Core

As shown by Freyer [45], cells in the inner regions of spheroids die due to low oxygen pressure, leading to the formation of a necrotic core surrounded by a viable rim of living cells. A central necrosis typically appears once spheroids reach diameters of approximately 500 − 800 *μm*. Although the thickness of the viable rim was once assumed to remain constant with spheroid growth, it is now recognized that in many cases the rim thins as the diameter increases [46]. Some models describing the development of this necrotic core assume that oxygen is completely consumed in the outer rim, imposing a zero-flux condition at the necrotic core-viable rim boundary [28,47]. Since, from a mechanistic standpoint, necrosis is primarily driven by oxygen depletion, we assume instead that cells die when the local oxygen concentration drops below a threshold value, while maintaining the zero-flux boundary condition at the tumorsphere center. Because CSCs exhibit greater plasticity to adapt to hypoxic conditions [48–51], we assign different hypoxic thresholds, *c*_*s*_and *c*_*D*_, to the CSC and DCC phenotypes, respectively, with *c*_*s*_ < *c*_*D*_. Tumorspheres are then grown as in previous cases, but all cells with oxygen concentrations below their respective thresholds are defined as *dead*. We also define the necrotic core radius, *R*_*n*_, as the maximum distance from the spheroid center at which viable DCCs are still present.

## 3. Results

In this section, we present the cell phenotype distributions in tumorspheres, both before and after the application of a single, uniform radiation dose. Successive applications are outside the scope of this study, as our goal is not therapy design but rather to assess the impact of spatial cell distributions. We therefore present histograms of cell spatial distributions, stratified by phenotype. As explained in Section 2 (Methods), the functions interpolating each histogram represent the probability density of finding a cell of a given phenotype at a distance *r* from the spheroid center. For comparison, in the following section, we also present the results of the therapy application to spheroids where the cells of both phenotypes are randomly distributed in their interior. CSC and DCC distributions will be labeled, respectively, *ρ*_*S*_and *ρ*_*D*_. In the last section, we examine the effect of radiation on tumorspheres with a necrotic core.

### 3.1. Phenotype distribution in a tumorsphere

Typical outcomes of the growth process in the absence of a necrotic core are shown in Fig.1 for *p*_*S*_ = 0.4, a value found compatible with the U87-MG cell line [32] and *p*_*S*_ = 0.7 (a CSC-enriched system) after simulating for 30 days (30 time steps), a relatively long period for a tumorsphere experiment. To better track the subpopulations present in the tumorsphere, proliferative CSCs are colored in red and proliferative DCCs in blue; the colors of the quiescent cells in each phenotype are displayed using lighter shades. The result for *p*_*S*_ = 0.4 is depicted in Fig. 1a, where a cross section of the tumorsphere shows that the CSCs are all quiescent, trapped near the center of the colony, and surrounded by DCCs. In contrast, for *p*_*S*_ = 0.7, Fig. 1b shows that there are CSCs in a proliferative state at the colony surface; they appear at the surface as red spots surrounded by proliferative DCCs (Fig. 1c). This is significant, given that many mathematical models in the literature assume a uniform CSC distribution. Furthermore, we carried out some preliminary experiments [22], confirming that the CSC distribution looks as predicted by the model, an agreement that highlights the power of simulations to guide experimental design.

**Figure 1:**
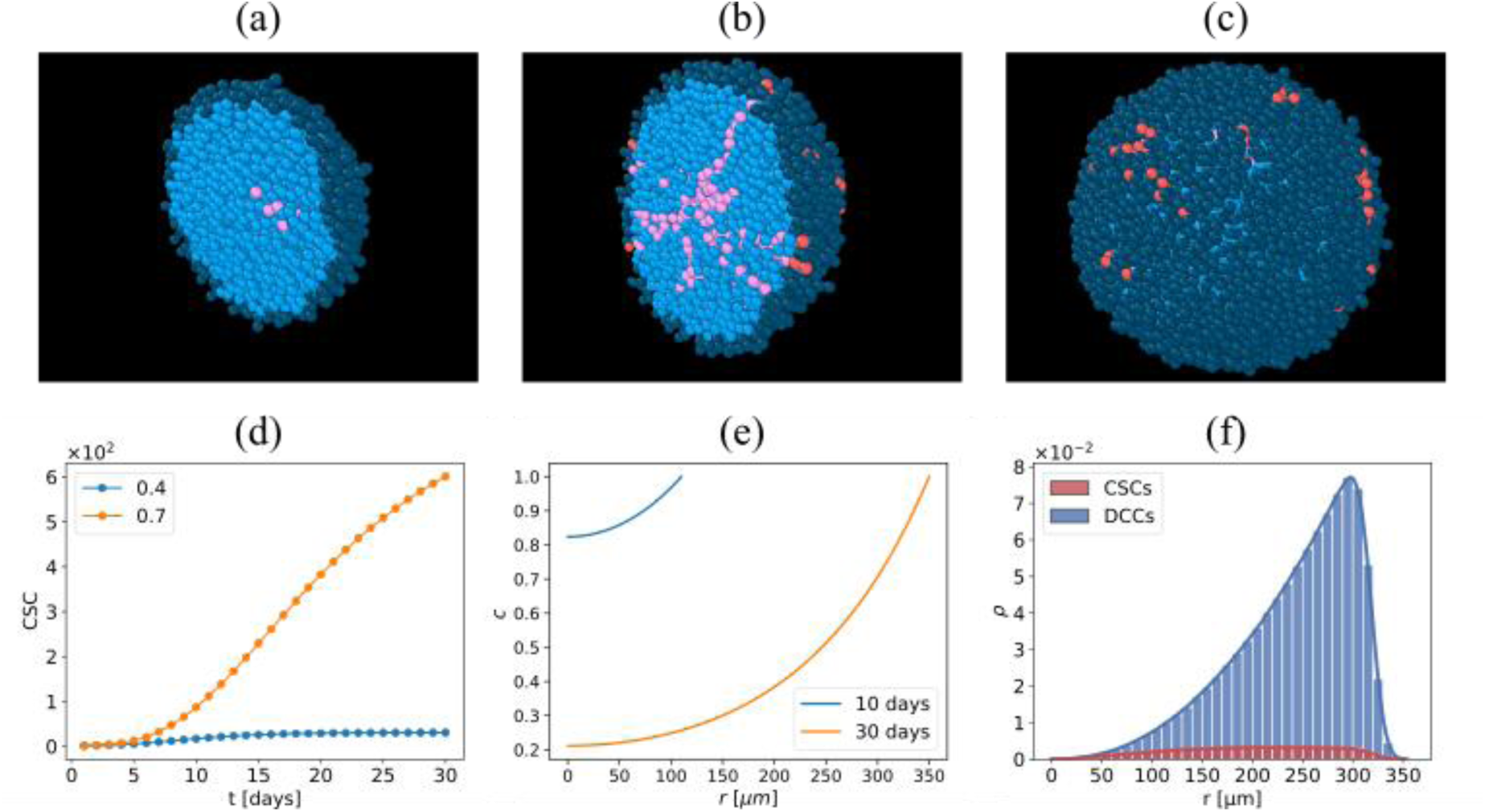
(a) Sagittal cross-section of a tumorsphere for self-renewal probability (*p*_*S*_ = 0.4). CSCs are quiescent and confined to the core. (b) Sagittal cross-section for a higher self-renewal probability (*p*_*S*_ = 0.7). CSCs can reach the periphery and keep proliferating. (c) External view of the tumorsphere for *p*_*S*_ = 0.7, showing proliferating CSCs (in red) at the surface. (d) Time evolution of CSC populations. For *p*_*S*_ = 0.4 (green) the growth of CSCs is arrested at an early stage. For *p*_*S*_ = 0.7 the population of CSCs keeps increasing but its growth rate decreases over time. (e) Oxygen concentration profile as a function of the distance r from the center for 10-day (black), 30-day (blue), and 60-day (orange) tumorspheres. (f) Total cell distribution inside a 30-day tumorsphere.

The evolution of the total number of CSCs is shown in Fig. 1d for an average over 32 realizations of the 30-day-old tumorspheres. As expected, the fastest growth in the CSC number occurs for the highest ps. Growth is arrested earlier for *p*_*S*_ = 0.4 (blue line), because CSCs become rapidly surrounded by DCCs and go to a quiescent state. For the higher self-renewal probability (*p*_*S*_ = 0.4, orange line), CSCs continue to proliferate for longer times, but their growth rate decreases monotonically. Ongoing physical analysis using percolation theory surprisingly indicates that, if the tumorsphere were allowed to grow indefinitely, at very long times, CSCs would appear at the surface only in the absence of differentiation (*p*_*S*_ = 0.4). Thus, for *p*_*S*_ < 1, it is unlikely, although theoretically possible, to find CSCs on the surface of large spheroids. Figure 1e shows the oxygen concentration profiles for 10-day-old (blue) and 30-day-old (orange) tumorspheres. The low oxygen concentration near the center of the bigger tumorspheres makes cells there especially radioresistant.

Because of the geometrical constraints, the tumorspheres have their cells roughly arranged in concentric layers. In Fig. 1f we depict the radial distribution of the cells of both phenotypes in 10 *μm*-thick shells for a 30-day-old tumorsphere. At the center of the tumorsphere, only the initial cell is present. As we move away from the center, the area of the layers increases as *r*^2^. As a consequence, more cells have room to proliferate and the proliferative phenotype predominates in the outer layers. In this region, the cells cannot fill the layers during the same timestep, and the number of cells per layer decreases. The CSC distribution is flattened and shifted toward the center because DCC creation and space restrictions hinder its growth, while the DCC distribution is skewed towards the periphery, with a higher concentration of cells near the surface.

### 3.2. Radiotherapy on Small Tumorspheres

In Fig. 2, we show histograms for 10-day-old tumorspheres. Prior to therapy, the average number of CSCs is approximately one-half that of DCCs (143 versus 319, respectively) and is slightly dominant near the tumorsphere center, while the DCCs are more abundant near the rim, as shown in panel (a). In panel (b), we show the surviving and killed cell distributions following therapy. Since the cells are well oxygenated everywhere in the spheroid, most cells of both phenotypes are killed and the protection offered by the remoteness from the surface is minimal: as a consequence of the therapy, the percentage of live CSCs had only a small increase: from 30.8% to 33.2%. The fraction of surviving CSCs (0.147) is barely higher than that of DCCs (0.135). The distributions of killed cells of both phenotypes are slightly biased towards the outer region.

**Figure 2:**
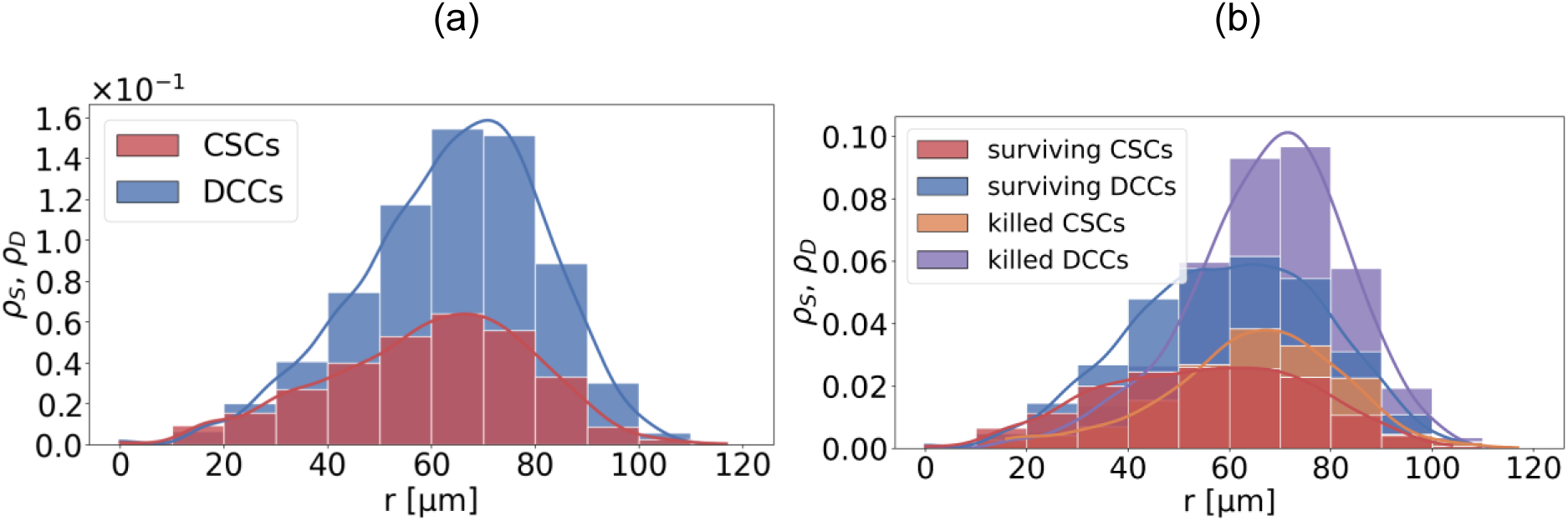
Normalized cell distributions for 10-day-old tumorspheres. (a) Before therapy, CSCs (red) and DCCs (blue) exhibit similar spatial distributions, although DCCs are slightly displaced outward from the center. (b) Following therapy, few cells of either phenotype remain, a consequence of the high oxygenation of the entire tumorsphere. The spatial distribution of killed cells closely resembles that observed prior to treatment.

### 3.3. 30-day Tumorspheres

We have shown that, in early-stage tumorspheres, spatial location is not important for CSC survival. The situation undergoes substantial changes when we consider bigger tumorspheres. Figure 3 shows histograms for 30-day-old tumorspheres immediately after therapy. The average number of CSCs before treatment was 1,999, from which 1,169 (58.5 %) survived, whereas 12,227 (48.7 %) from the original 25,089 DCCs survived. The CSC fraction of the total cell number, which was 0.074 before the therapy, grew to 0.088 afterward (see Table 1). Figure 3a displays the distribution of cells of each phenotype as a function of the distance from the tumorsphere center. A large fraction of the DCCs were killed near the surface, leading to a truncation of the peak in the surviving DCCs distribution. Because the number of DCCs is much larger than that of CSCs, the CSCs distributions are magnified in Fig. 3b. The black line represents the kernel density estimate of the total CSC population, which has a relatively flat profile and drops off at *r* = 290 *μm*, close to the maximum of the DCCs distribution. After therapy, the total CSC distribution separates into two well-defined peaks.

**Figure 3:**
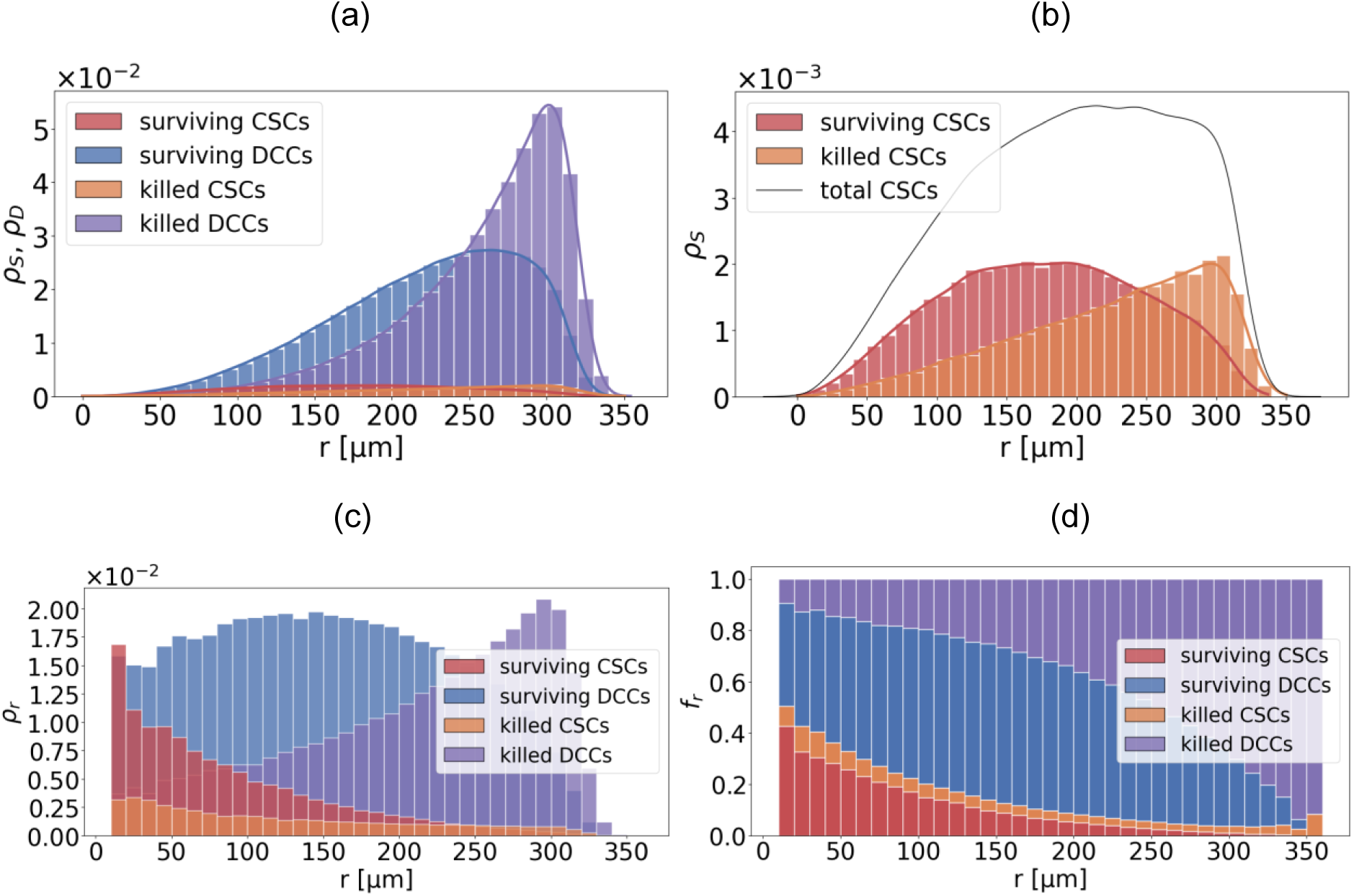
Cell distributions for 30-day-old tumorspheres after therapy. (a) Histograms of killed and surviving CSCs and DCCs. For both phenotypes, cells were preferentially killed near the surface. (b) Expanded view of the distributions of surviving and killed CSCs. (c) Radial cell density shows CSCs are mainly located far from the surface. (d) Radial dependence of cell fractions reveals enhanced protection in the core of the tumorsphere.

**Table 1:**
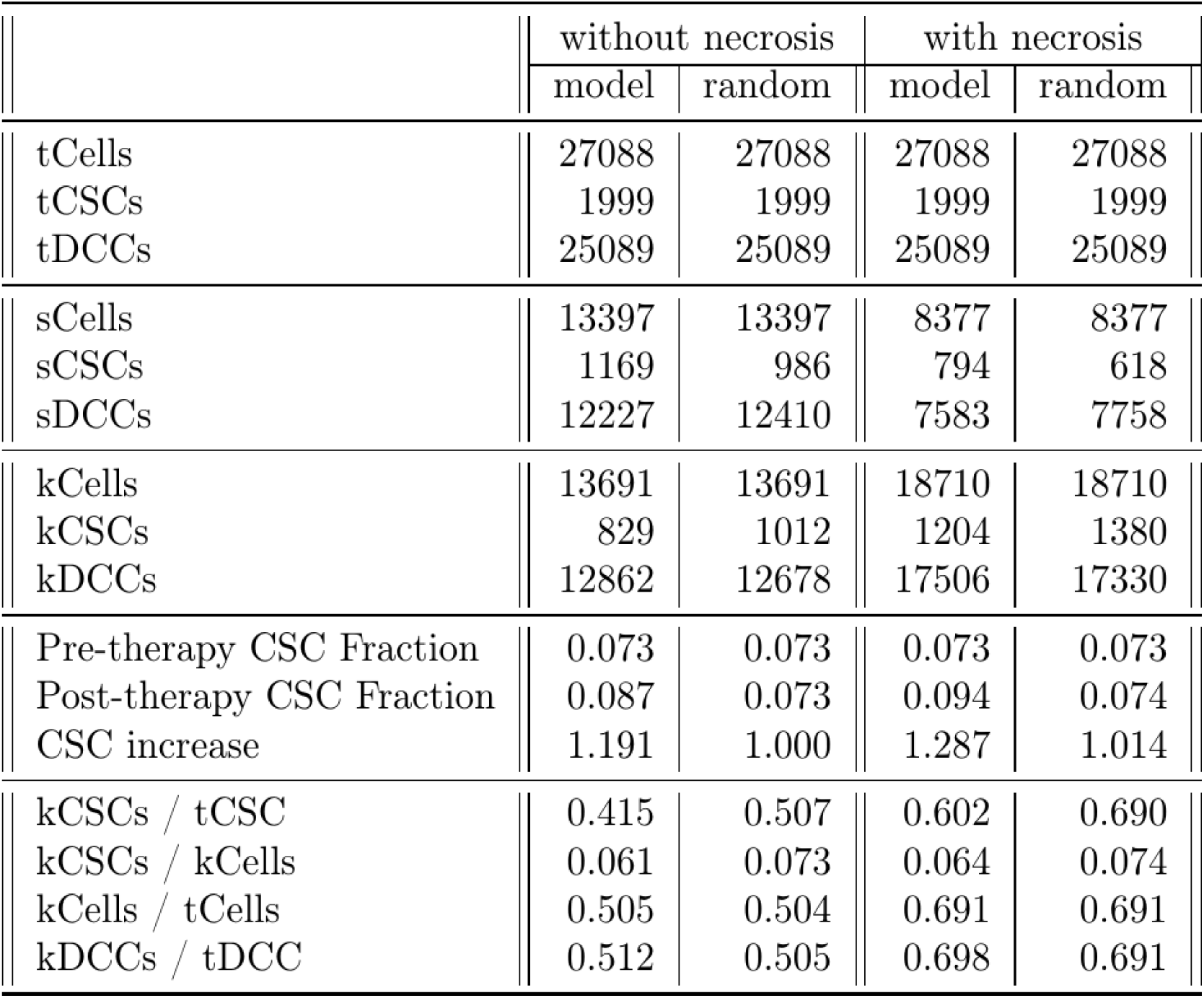
Cell population statistics for *p*_*S*_=0.7. The total population (t) of cells is broken down by phenotype (CSCs and DCCs) and by killed (k) and surviving (s) cells.

The peak of the surviving CSC distribution (red) shifts toward the center while maintaining a relatively flat profile, whereas the peak of the killed CSC distribution (orange) shifts toward the surface. The latter exhibits an approximately linear increase with the radius in the region to the left of the maximum. A comparison of panels (a) and (b) shows that, while the maxima of the surviving cell distributions are well separated, those of the killed cell distributions nearly coincide. A different perspective emerges if we look at the radial dependence of the cell densities depicted in Fig. 3c. For both killed and surviving CSCs, the density decreases with increasing radius, reflecting their abundance in the tumorsphere core. In contrast, the density of killed DCCS (violet) increases towards the surface, consistent with their pre-therapy DCC predominance in that region. The protective nature of the inner regions is further evidenced by the radial density of surviving DCCs (blue), which has a broad maximum far from the surface. For clarity, Fig. 3d shows the fractions of each type of cell as a function of the distance from the center. The fraction of surviving cells for both phenotypes is similar near the center, but decays with radius for the CSCs. In the outer rim, the fraction of killed cells approaches unity, with most of them being DCCs. This last panel reveals the role of the self-renewal/differentiation mechanism in enhancing the effective radioresistance of CSCs.

### 3.4. Comparison with Random CSC Distributions

To reinforce our conclusions on the impact of the non-uniformity of the CSC distribution on their survival, we implement the same radiotherapy dose to spheroids obtained from the tumorspheres studied in section 3.3 by redistributing both cell phenotypes at random, as explained in the Methods section. The results are depicted in Fig. 4, whose panel (a) shows the distributions of surviving and killed CSCs immediately after a therapy application at *t* = 30 days. The histograms for the CSCs and DCCs exhibit the same spatial profiles, albeit at different scales, as seen by comparing with panel (b), which displays the CSC distributions alone. In this panel, we compare the distributions from our model to those resulting from a random distribution of CSCs immediately before the therapy. When the pre-therapy distribution is generated by our model, surviving CSCs are positioned closer to the tumorsphere center than their randomly generated counterparts. They are also noticeably more numerous than those that originated from the random distribution.

**Figure 4:**
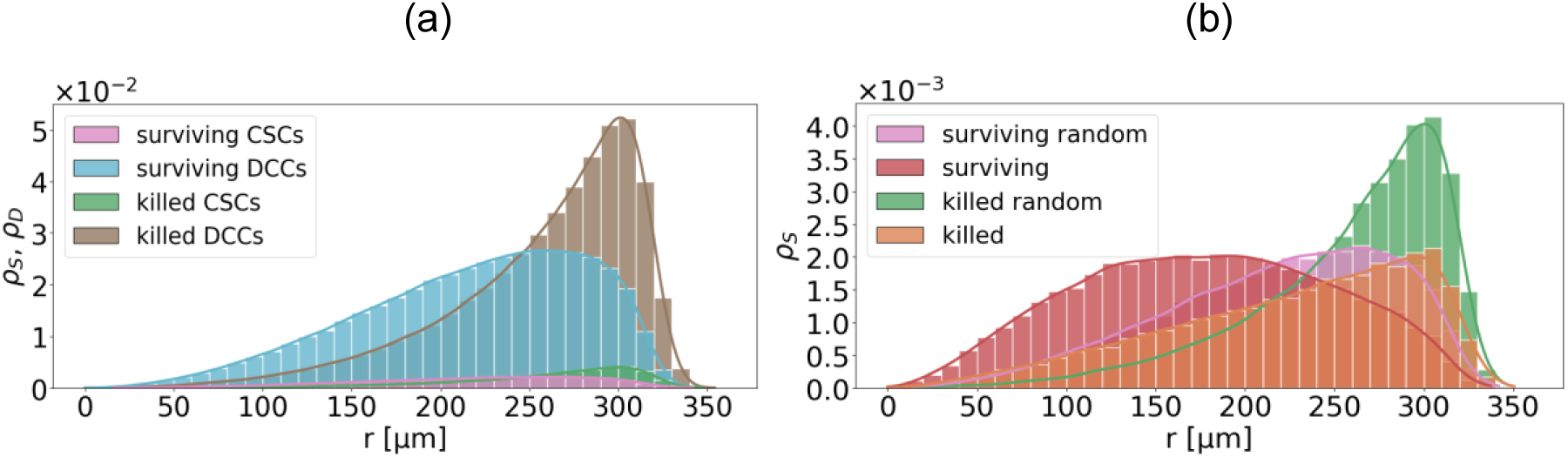
Effect of radiotherapy on a 30-day-old tumorsphere with a random CSC distribution. Histograms show cell numbers as a function of distance from the tumorsphere center. (a) CSC and DCC spatial distributions display the same overall shape, although their total numbers differ greatly. (b) Comparison between the CSC distributions in our model and those resulting from a random cell allocation. The number of surviving CSCs is higher in our model.

A complete set of statistical values is reported in Table 1. These results clearly show that the killed CSC fraction was lower in our model than in the random distribution (41.5% vs. 50.7%), while the number of surviving CSCs was 19% higher. The reason for this difference is that more cells get killed near the rim, where the *O_2_* concentration is high, and where our model predicts fewer CSCs to be located.

### 3.5. 30-day-old tumorspheres with a necrotic core

Large tumorspheres typically develop a necrotic core, making it important to assess how such a core influences the results presented in the previous sections. We do this for 30-day-old simulated tumorspheres, which reach a typical radius of *R* = 350 *μm*. The size of the necrotic core depends on various parameters. Here we use the data from Ref. [52], which reports, for an MCF-7 spheroid with *R* = 350 *μm*, a core radius of *R*_*n*_ = 150 *μm*. We must estimate the values for the cell death thresholds, *c*_*D*_ and *c*_*S*_, which satisfy *c*_*D*_ > *c*_*S*_. From Eq. (1), under our boundary conditions, we obtain *c*(*r* = 150 *μm*) = 0.3. Thus, we set *c*_*D*_ = 0.3, and, since CSC oxygen consumption varies widely [17], depending on factors such as the cell line, and given the lack of reliable data on the CSC fraction, we choose, without loss of generality, *c*_*S*_ = 0.23. To our current knowledge, there are no reports on the fraction of CSCs that survive under anoxic conditions, if any. Thus, the chosen values assume there is a non-negligible CSC fraction that dies from anoxia.

By definition, *R*_*n*_ is the maximum radius within which there are no live DCCs; because *c*_*D*_ > *c*_*S*_, a small number of CSCs may still survive within the core, but their oxygen consumption is negligible, so we may calculate the oxygen distribution disregarding this small population, i.e., *c*(*r*) = 0 for *r* < *R*_*n*_. Under these conditions, we simulated tumorsphere growth and applied a single radiotherapy dose on day 30, as in previous cases. The cell distribution immediately before therapy is shown in Fig 5a, where the necrotic core has reached a radius of *R*_*n*_ = 150 *μm*, visible as the transition between dead (cyan) and live (blue) DCCs. CSCs survive hypoxia within the core, down to about 70 *μm* (see inset for a magnified view of this region). Varying *c*_*S*_merely shifts the boundary between dead and live CSCs. The chosen value of *c*_*S*_is very conservative as it results in the killing of many CSCs near the center, leading to their complete annihilation in large tumorspheres (sup Fig. S4). As a consequence, our simulated tumorspheres represent the worst possible case. Figure 5b shows that therapy produces the same distribution profiles as those observed in the case without a necrotic core, cf. Fig. 3, except for the truncations due to necrosis. The number of therapy-killed DCCs and CSCs remains in the same proportions as in Fig. 3c. As in Fig. 3b, live CSCs outnumber dead CSCs near the center (the opposite occurs for DCCs). The convex profile of killed DCCs and the concave profile of live DCCs are likewise preserved.

**Figure 5:**
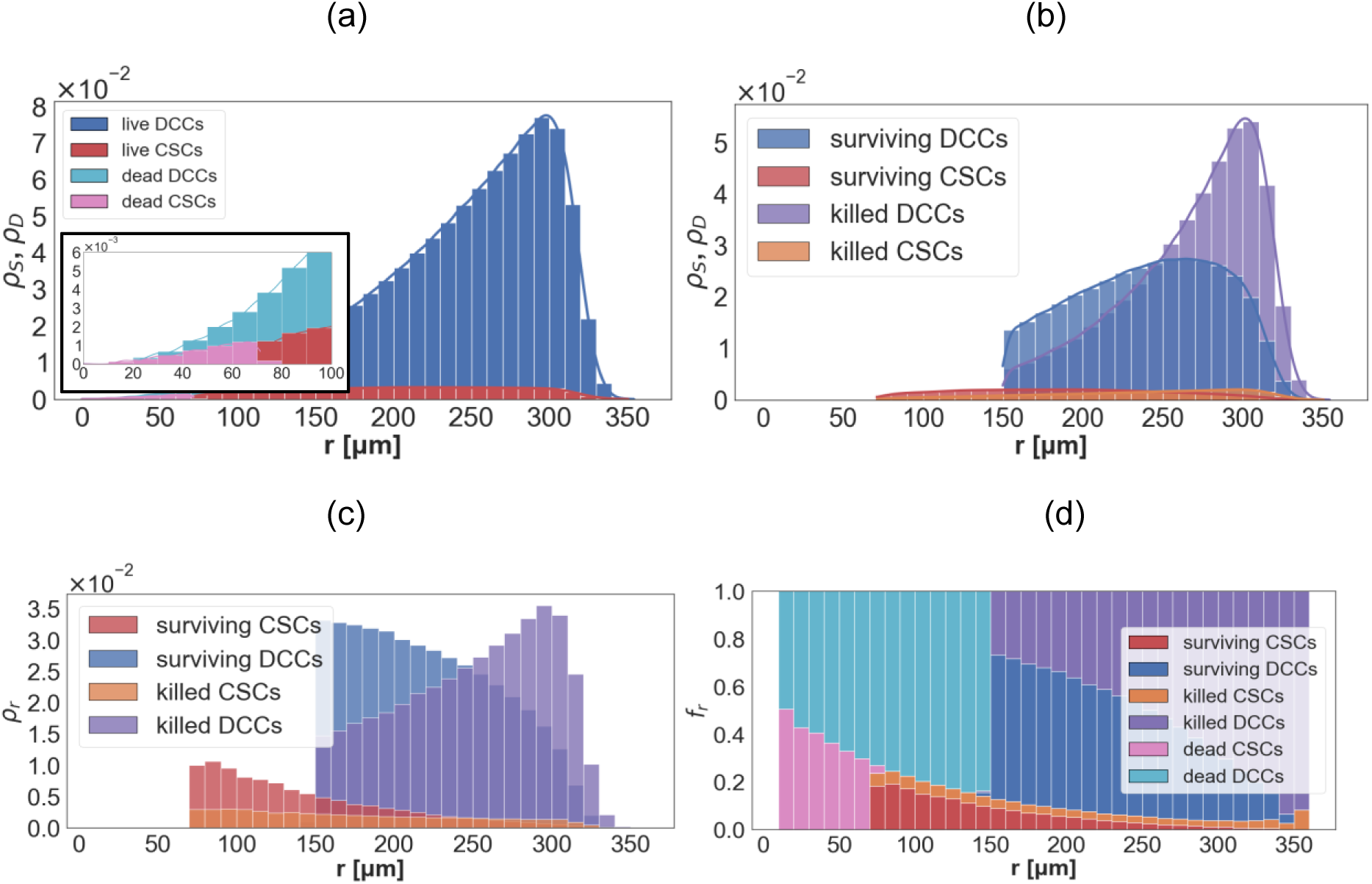
Combined effects of hypoxia and radiation on 30-day tumorspheres. (a) Cell distribution before the therapy. The spheroid has developed a necrotic core of radius *R*_*n*_ = 150 *μm*. The inset highlights the hypoxic CSC range. (b) Cell distribution after a single dose of radiotherapy; the fraction of killed DCCs in the necrotic core exceeds that of killed CSCs. (c) Histograms from (b) rescaled to show the radial dependence of the cell concentration. (d) Concentration fractions of all cell types. Here *c*_*S*_ = 0.23 and *c*_*D*_ = 0.30.

As shown in Table 1, the average number of CSCs before treatment was 1,999, of which 497 were already in the necrotic core at the time of the irradiation. Subsequently, 707 CSCs were killed by radiation, leaving 794 survivors that represent 39.7% of the original CSC population. Among the 25,089 DCCs present at the time of treatment, 6,780 were already necrotized by hypoxia; and 10,725 were subsequently killed by radiation. The remaining 7,583 DCCs represent 30.2% of the original DCC population, a significantly lower survival fraction than that of CSCs. The CSC fraction of the total cell population, which would have been 0.073 without hypoxia, rose to 0.076 once the effects of hypoxia were taken into account, and finally increased to 0.094 following therapy, higher than the value of 0.087 predicted in the absence of necrosis (see Table 1).

To better understand the interplay between radiotherapy and the necrotic core, we divided the number of cells in each radial bin by the area of the corresponding spherical shell to obtain the radial cell density distribution. Figure 5c shows that a higher density of CSCs survives near the tumorsphere center, whereas most DCCs are eliminated by therapy in its periphery. The concentration fraction of each phenotype is presented in Fig. 5d, with each bar representing the relative abundance of each cell phenotype in a given shell. The inner necrotic core is fully devoid of live cells, while the outer necrotic core contains about 20% of the CSCs that survived both hypoxia and radiation. In the viable rim (*r* > *R*_*n*_), DCCs, both dead and alive, are the dominant phenotype, and the CSCs fraction is minimal.

Next, we compare our results with those obtained under the assumption that CSCs and DCCs are randomly distributed throughout the tumorsphere. In this case, only 618 CSCs survive (see Table 1), a substantially lower number than the 794 survivors predicted by our model with spatial structure. Moreover, in the random distribution case, the ratio of the surviving CSCs to the total cell population (0.074) remains unchanged by either hypoxia or radiation.

In the Supplementary Information, we present results for a 60-day tumorsphere. Although this scenario is not fully realistic, since additional processes such as cell senescence or the accumulation of toxic metabolites would likely intervene, it illustrates the expected fate of the spheroid under continued growth. Except deep in the spheroid interior, the population of CSCs becomes minimal. The emergence of a large necrotic core eliminates most of these cells, leaving a viable outer rim composed almost solely of DCCs.

## 4. Discussion

Cancer stem cells are widely reported to exhibit enhanced survival following radiotherapy, a phenomenon commonly attributed to intrinsic radioresistance. In this work, we show that spatial organization alone can generate a substantial survival advantage for CSCs, even in the absence of intrinsic differences in radiosensitivity. This increased CSC survivability can be explained by their preferential location in the interior of the tumorsphere, where oxygen levels are lower and radiation-induced cell killing is reduced.

To support this hypothesis, we first determined the location of the CSCs within a tumorsphere, which we achieved by extending a growth model first proposed in [23], whose predictions were corroborated by the experimental observations of Ref. [22]. As shown in Fig. 1, CSC lineages create paths in the tumorsphere bulk that are eventually constrained by spatial limitations that force them into quiescence. As a consequence, most CSCs reside well in the tumorsphere interior. This agrees with the observation by Li and co-workers of CSCs in regions near necroses, which led them to suggest the existence of a hypoxic niche for the CSCs [53].

To focus on the relationship between CSC positioning and survival, we deliberately ignored possible intrinsic radiosensitivity differences between CSCs and DCCs and excluded complicating, but non-essential processes, such as cell shedding [54], senescence [32], and the secretion of growth-limiting factors. Since our objective is not to model cancer growth, but to analyze the relation between CSC location and survival, we chose the simplest possible description. Radiotherapy was modeled as a single, spatially uniform dose, with a survival probability that depends linearly on the local oxygen concentration, *S*(*r*) = 1 − *c*(*r*), normalized such that *c*(*R*) = 1. This is a conservative assumption. In real systems, there is typically an oxygen concentration threshold above which cells are close to maximally radiosensitive. As a result, our model likely underestimates cell killing in the well-oxygenated periphery -away from the CSCs- and therefore underestimates the fraction of CSCs in the surviving population (a biologically accurate *S*(*r*) *w*ould amplify the effect). Even with our assumption of a linear decrease in cell elimination with decreasing oxygen concentration, we showed that the fraction of surviving CSCs is markedly higher than that of DCCs. Our formulation isolates the contribution of spatial oxygen gradients, independent of intrinsic radiosensitivity differences between CSCs and DCCs.

Our results were further confirmed by simulating the application of the same radiotherapeutic dose to tumorspheres obtained by randomly redistributing the CSCs and DCCs, both with and without necrosis. These simulations showed that:

i. the surviving CSC fraction is considerably higher in the tumorspheres generated by the growth model;
ii. the difference between these survival fractions increases with time, i.e., with tumorsphere size;
iii. the surviving DCC fraction is slightly higher for the random tumorspheres, as these contain more DCCs in the hypoxic interior, and
iv. in the random tumorspheres, the surviving CSC and DCC fractions are nearly identical, confirming that the survival advantage in our model is a property of spatial emergence rather than cell-type identity. This is a strong validation of our hypothesis.

Together, these results support the conclusion that CSC spatial positioning is an important determinant of radioresistance.

A secondary mechanism that may further enhance CSC survival is quiescence. As the spheroid grows, central cells become quiescent and, consequently, consume less oxygen, enhancing radiation resistance. Since CSCs are predominantly positioned far from the surface, quiescence amplifies their chances of surviving the therapy. Because of its structure, our model can also identify which cells become quiescent as the tumorsphere grows. A quantitative assessment of the contribution of quiescence to CSC survival is an aspect that can be addressed within the present framework.

Since large tumorspheres usually develop a necrotic core, we also modeled the CSC-driven growth of a tumorsphere under hypoxic conditions, where cells far from the surface die from oxygen deprivation. Due to their greater adaptability to low oxygen levels, some CSCs were allowed to survive within the core. Simulation results indicate that the radiotherapy effects are similar to those observed in non-necrotic tumorspheres. Necrosis is associated with the development of invasion and metastasis [55]. The survival of CSCs in necrotic regions suggests that the boundary of necrotic regions may provide a microenvironment favorable to plasticity and dedifferentiation. The de-differentiated phenotype has been reported to be particularly common in metastatic tumors [7]. These findings are consistent with the idea that microenvironmental conditions can shape CSC behavior and, ultimately, tumor aggressiveness.

Although our analysis does not focus on dose (*D*) dependence, the framework can be extended to incorporate standard radiobiological models such as the Linear-Quadratic (LQ) formalism. Notably, the survival fraction, *SF*(*D*), predicted by the LQ model does not incorporate spatial information [29, 56, 57]. In this context, the overall survival fraction could be approximated by combining the spatially resolved survival profile obtained here with a dose-dependent survival function *SF*(*D*). This approach implicitly assumes a separable contribution of spatial oxygen effects and dose dependence, which is a simplification of the full radiobiological response.

To test our hypothesis on the role of CSC positioning in radioresistance, we modeled tumorsphere growth as a simplified system initiated from a single cancer stem cell. Although idealized, this approach indicates how spatial organization can affect treatment outcomes, offering insights that may generalize to more complex neoplastic systems. In particular, the observed link between CSC location and survivability could be exploited to optimize radiotherapy, whether applied alone or in combination with other treatments. Of course, translating these findings to real tumors will require incorporating additional biological processes such as cell migration [58], plasticity [59–61], and senescence [32, 62]. Nevertheless, our results underscore the importance of spatial heterogeneity [63, 64] in shaping therapy response. Recently, Syga et al. developed a cellular automaton model that they used to examine the effects of phenotypic switching between migration and proliferation, showing that higher phenotypic heterogeneity is associated with poorer therapeutic outcomes [65]. In this work, we have shown that the spatial localization of this heterogeneity further exacerbates the problem. In this context, we also note a recent study that uses a multiscale mathematical model to investigate how the tumor microenvironment influences acquired drug resistance, with a particular focus on drug-tolerant persister cells [66].

In summary, our results indicate that the enhanced survival of CSCs following radiotherapy need not arise solely from intrinsic radioresistance. Instead, it can emerge naturally from the interplay between spatial organization and oxygen-dependent radiation response. This mechanism suggests that therapeutic strategies targeting tumor spatial structure or oxygenation may complement conventional approaches aimed at intrinsic cellular sensitivity.

## Acknowledgments

This work was supported by SECyT-UNC (Consolidar project 33620230100392CB) and CONICET (PIP 11220200103005CO). We are grateful to Luciano Vellón and Ariel Martínez for illuminating discussions. This work used computational resources from UNC Supercómputo (CCAD) Universidad Nacional de Córdoba (https://-supercomputo.unc.edu.ar), which are part of SNCAD, República Argentina.

## Captions – Supporting Information

**Figure S1:** *UML* diagram of the simulation package. Simulation aggregates Culture objects, while Culture is composed of Cell objects. Output classes implement the TumorsphereOutput interface, and SpatialHashGrid is used by Culture for efficient neighborhood searches.

**Figure S2:** *Cell distributions for 60-day-old tumorspheres after therapy.* (a) Killed and surviving CSC and DCC distributions display the same shape as those for 30-day old tumorspheres. The CSC population is too small to be clearly visible and is therefore rescaled in the next panel. (b) Expanded view of the surviving and killed CSC distributions. (c) Radial cell density shows CSCs are mainly located far from the surface. (d) Cell fraction profiles reveal that cells would be protected within the core of the tumorsphere. No killed CSCs are detectable at this scale.

**Figure S3:** *Effect of radiotherapy on a 60-day-old tumorsphere with a random CSC distribution.* Histograms show cell numbers as a function of distance from the tumorsphere center. (a) CSC and DCC spatial distributions display the same overall shape, although their total numbers differ greatly. (b) Comparison between the CSC distributions in our model (cf. Fig S2b) with those resulting from a random cell allocation shows that the CSC survival is higher in our model

**Figure S4:** *Combined effects of hypoxia and radiation on 60-day tumorspheres* for *c*_*S*_ = 0.23 and *c*_*D*_ = 0.30. (a) Prior to therapy, the tumorsphere has developed a necrotic core of radius *R*_*n*_ = 590 *μm*. The inset highlights the hypoxic CSC range, showing a very small fraction of live CSCs located far from the center due to the stringent threshold. (b) Cell distribution after a single dose of radiotherapy; for clarity, cells that were already dead at the time of the therapy are not shown. (c) Histograms from (b), rescaled to show the radial dependence of the cell concentrations. (d) Cell fraction profiles for all cell types, highlighting the CSC annihilation resulting from the strict anoxic threshold *c*_*S*_. Insets show enlarged views of the region containing CSCs.

**Table S1:** Cell population statistics for the cases studied (*p*_*S*_ = 0.7). The total population (*t*) of cells is broken down by phenotype (CSCs and DCCs) and by cell fate: killed (*k*) and surviving (s) cells.

